# Shared and divergent features of the immune response to acute and chronic *Trypanosoma cruzi* infection in mice revealed by cardiac transcriptomics

**DOI:** 10.64898/2026.11.02.763600

**Authors:** Francisco Olmo, Shiromani Jayawardhana, Martin C. Taylor, John M. Kelly, Amanda Fortes Francisco

## Abstract

Chagas disease is caused by infection with *Trypanosoma cruzi* and can result in chronic cardiomyopathy. However, the molecular programmes that define infected cardiac tissue, and how they change as the host-parasite relationship moves from acute pathogenic infection to long-term persistence, remain incompletely understood. Here, we used RNA sequencing to characterise host transcriptional responses in cardiac tissue during both acute and chronic infection in a mouse model. Cardiac parasite distribution was defined using bioluminescence imaging and histology, and RNA sequencing was performed on anatomically standardised cardiac tissue samples collected at both stages. Acute infection was associated with a broad interferon-driven immune activation, whereas during chronic infection this network contracted, with preservation of a tightly connected antigen presentation core centred on major histocompatibility complex genes and immunoproteasome components. Genes associated with extracellular matrix remodelling and fibrosis were not broadly activated. These findings indicate that reduced, but sustained immune activation, rather than widespread fibrotic transcriptional remodelling, represents a prominent feature of cardiac tissue in chronically infected mice. They identify a transition from an overtly inflammatory, pathogen-driven acute response to a more restricted chronic host-parasite equilibrium in which immune surveillance remains transcriptionally active despite very low and spatially intermittent cardiac infection. We identified a conserved 20-gene signature shared between acute and chronic stages. This overlap is consistent with continued or episodic parasite-derived stimulation and/or intrinsic host mechanisms that maintain cardiac immune surveillance. Thus, chronic *T. cruzi* infection can be viewed as a persistent host-parasite association in which biological coexistence does not imply benignity: immune containment and parasite persistence can occur alongside cumulative cardiac injury.

## 1 Introduction

Chagas disease is a major public health challenge across the American continent and is increasingly recognized globally due to migration and non-vectorial transmission routes such as blood transfusion and congenital infection (de Sousa et al., 2024). Affecting an estimated 10.5 million people, Chagas disease is the leading cause of infectious myocarditis in Latin America and contributes substantially to morbidity and mortality (Pérez-Molina and Molina, 2018; GBD 2023 Chagas Disease and RAISE Study Collaborators, 2026). A hallmark of infection with the causative agent *Trypanosoma cruzi,* is its broad tissue tropism: the parasite can infect virtually any nucleated cell, with cardiac, gastrointestinal, adipose, skeletal muscle and skin tissues representing important sites of infection (Lewis and Kelly, 2016; Lewis et al., 2016; Ward et al., 2020; Ward et al., 2020; Bonney and Engman, 2008; Pérez-Mazliah et al., 2021). The absence of non-invasive technology that facilitates monitoring of real-time parasite dissemination in humans has been a limiting factor in studying disease progression.

The clinical course of Chagas disease is complex and often silent. The initial acute phase is typically mild or asymptomatic, although myocarditis develops in some cases. The subsequent chronic phase of infection can persist for decades, with about 30-40% of individuals eventually developing cardiac disease. Transition to life-threatening pathology has been closely linked to parasite persistence and the host immune response (Lewis and Kelly, 2016; Bonney and Engman, 2008; Pérez-Mazliah et al., 2021). In the heart, infection drives myocarditis, tissue remodelling and fibrosis. However, the complex long-term nature of Chagas disease in humans has restricted research on disease pathogenesis, and the spatiotemporal dynamics of parasite localization and host-parasite interactions remain poorly resolved. Thus, an important unresolved question is how a persistent host response is maintained when parasite burden within the heart itself is low or intermittent, and how this relationship contributes to progressive cardiac disease.

Mouse models provide an opportunity to investigate this relationship experimentally (Khan et al., 2024). Although RNA sequencing (RNA-seq) studies have characterised transcriptional responses in *T. cruzi*-infected murine hearts (de Castro et al., 2020; Navarro et al., 2015; Goldenberg et al., 2009; Ferreira et al., 2017), less is known about how these responses change between acute and chronic infection when considered alongside the marked temporal and spatial changes in parasite burden and distribution. This distinction is particularly important because cardiac pathology may reflect spatially heterogeneous and cumulative processes. A key obstacle lies in the inability to sensitively and specifically track *T. cruzi* infection in vivo over time and across tissue compartments. Conventional histopathology, although definitive, is terminal and inherently limited by sampling bias and lack of spatiotemporal resolution (Lewis et al., 2014).

Bioluminescence imaging (BLI) provides an approach to address this problem by allowing parasite distribution to be monitored longitudinally and with high sensitivity (Costa et al., 2018; Taylor et al., 2021; Taylor et al., 2020; Olmo et al., 2024). Reporter parasite lines have revealed marked changes in tissue distribution between acute and chronic infection and have identified the gastrointestinal tract and skin as important sites of persistence during chronic murine infection (Ward et al., 2020; Ward et al., 2020). Combining information on parasite distribution with anatomically standardised cardiac transcriptomics therefore provides an opportunity to distinguish transcriptional responses associated with widespread acute cardiac infection from those maintained during chronic infection when cardiac parasite burden is substantially reduced and more localized.

Transcriptomic studies have been used to advance our understanding of host responses during *T. cruzi* infection. RNA-seq of infected hearts has revealed immune-metabolic remodelling, inflammatory activation, and strain-specific gene expression (de Castro et al., 2020; Navarro et al., 2015; Goldenberg et al., 2009; Ferreira et al., 2017). However, in the absence of information on the sites where parasites reside in the myocardium, it has not been possible to determine whether transcriptional changes reflect local parasite proximity, systemic inflammation or disease stage. For example, imaging studies show that during chronic infections, parasites persist at very low numbers in discrete, dynamic foci, often outside the heart, even as cardiac pathology progresses (Lewis et al., 2016; Ward et al., 2020; Ward et al., 2020).

Here, we combined BLI of *T. cruzi* distribution with RNA-seq of anatomically standardised mid-right ventricular myocardial biopsies to define cardiac host responses during acute and chronic infection. We asked whether the marked reduction in cardiac parasite burden accompanying transition to chronic infection is associated with resolution of the acute transcriptional response or with persistence of a defined host-response programme. Comparison of the two disease stages identified a conserved 20-gene signature dominated by antigen presentation, interferon (IFN)-associated responses and immunoproteasome components, despite major differences in parasite burden and cardiac infection dynamics. These findings provide a framework for understanding how immune surveillance can persist and evolve within cardiac tissue during chronic infection, when local parasite burden is markedly reduced and cardiac infection is intermittent.

## 2 Material and methods

### 2.1 Ethics Statement, animals and infection

All animal experiments were conducted under UK Home Office project license PPL P9AEE04E4 and approved by the London School of Hygiene and Tropical Medicine Animal Welfare and Ethical Review Board. All procedures complied with the UK Animals (Scientific Procedures) Act 1986. BALB/c mice were purchased from Charles River (UK). Upon arrival, they were randomly assigned between cages and acclimatized for at least 5 days. Animals had free access to food and water and were maintained under a 12 h light/dark cycle at controlled humidity and temperature. Mice were monitored daily for signs of clinical disease.

### 2.2 Parasite strains, infection, and histology

Two transgenic derivatives *T. cruzi* CL Brener that stably express the red-shifted luciferase PpyRE9h were used: the bioluminescent CL-Luc line (Lewis et al., 2014) and the dual bioluminescent-fluorescent CL-Luc::mScarlet line (Costa et al., 2018; Taylor et al., 2021). The CL-Luc::mScarlet line enables spatial visualization and histological confirmation of parasite persistence in tissues, at single cell resolution (Ward et al., 2020; Ward et al., 2020). Bloodstream trypomastigotes were purified and quantified as previously described (Lewis et al., 2014). Female BALB/c mice, aged 7-8 weeks, were infected intraperitoneally with 1×10³ trypomastigotes.

For histological analysis of parasite location within cardiac tissue during the acute phase, an independent cohort of BALB/c mice (n=5) was infected. Hearts were collected at 14 days post-infection (dpi), fixed in 95% ethanol, embedded in paraffin, and sectioned according to established protocols (Ward et al., 2020; Ward et al., 2020). Sections were mounted on glass slides, deparaffinized, and counterstained with 4′,6-diamidino-2-phenylindole (DAPI). Parasites expressing mScarlet were visualized by fluorescence microscopy, enabling identification of intracellular amastigote nests within cardiac tissue.

### 2.3 In vivo and ex vivo bioluminescence imaging

Whole-body bioluminescence imaging was performed using the IVIS Spectrum (Revvity, Hopkinton, MA, USA). For in vivo imaging, mice were injected intraperitoneally with D-luciferin potassium salt (200 mg/kg in sterile D-PBS), anesthetized with 2.5% isoflurane in oxygen, and imaged in the ventral and dorsal position using Living Image v4.8.2 (Lewis et al., 2014). For ex vivo imaging (Lewis et al., 2016; Lewis et al., 2014), mice were similarly injected with D-luciferin and euthanized 5 min later by exsanguination via the axillary vein under terminal anaesthesia (dolethal solution from Vetoquinol at 200 mg/ml 150 µl i.p.). Following removal of residual blood, organs and tissues, the remaining carcasses were placed in a square Petri dish containing D-luciferin solution (0.3 mg/ml in D-PBS) and imaged immediately.

### 2.4 Tissue collection, RNA extraction and sequencing

For RNA-seq, a fresh 3 mm mid-right ventricular myocardium biopsy was collected from BALB/c mice at both acute and chronic stages of infection. The cardiac punch biopsy was collected from the same area from all hearts to ensure reproducible anatomical sampling across experimental groups and to avoid selecting biopsy sites on the basis of local BLI intensity. For the acute stage, heart samples were obtained from uninfected control (n=5) and infected (n=10) mice at 11 days post-infection (dpi), corresponding to the early acute phase of infection. For the chronic stage, the same number of BALB/c mice were sampled at 171 dpi, corresponding to the late chronic phase of infection. These time points were deliberately selected to capture early cardiac transcriptional responses during the acute phase and later host transcriptional responses during chronic infection. All cardiac biopsies were added to TRIzol reagent (Invitrogen), with RNA extraction on the same day using the RNeasy Mini Kit (Qiagen). RNA purity and concentration were measured using NanoDrop and Qubit 2.0, and RNA integrity was assessed with an Agilent 2100 Bioanalyzer.

### 2.5 Library preparation and sequencing

Library preparation, sequencing, and primary bioinformatic processing were performed by BMKGene (BMKCloud platform), with downstream analyses and interpretation conducted by the corresponding author. mRNA was enriched using oligo(dT)-attached magnetic beads, randomly fragmented, and reverse transcribed into cDNA using random hexamer primers. Second-strand synthesis was performed with DNA polymerase I, followed by end repair, A-tailing, adaptor ligation, and PCR amplification. Libraries were purified with AMPure XP beads, and quality was assessed using Qubit 2.0, Agilent 2100 Bioanalyzer, and qPCR. For the acute and chronic studies, across the 15 libraries for each disease stage, a total of 192.7 Gb and 200.3 Gb of clean data were generated, respectively, with at least 12.2 Gb per individual sample. Sequencing quality was high, with >98.8% of clean bases at Q30 or higher, implying a Phred quality score of 30, which corresponds to a base-calling error rate of 0.1% (99.9% accuracy), a standard benchmark for high-quality Illumina sequencing.

RNA-seq libraries for acute and chronic infection stages were prepared and sequenced independently due to the temporal separation between experiments. Differential expression analysis was performed separately for each stage by comparing individual infected versus individual uninfected control hearts processed under the same experimental conditions. All downstream analyses were conducted independently for each stage using identical statistical thresholds to enable qualitative comparison of stage-specific transcriptional programmes. Accordingly, direct statistical comparison of expression levels between the acute and chronic sequencing experiments was not performed.

### 2.6 Data processing and alignment

Raw reads were processed to remove adapters and low-quality bases. Sequencing randomness, insert size distribution, and library complexity were examined to ensure quality. Clean reads were aligned to the *Mus musculus* reference genome (GRCm39, Ensembl release 104) using HISAT2 v2.2.1, achieving mapping rates of 98.0-98.5% across samples.

### 2.7 Expression quantification and differential analysis

Gene expression levels were quantified using StringTie (v2.1.4), and transcript abundances were reported as fragments per kilobase of transcript per million mapped reads (FPKM). To reduce background noise and exclude low-abundance transcripts, genes with expression levels ≤5 FPKM across all samples were removed prior to downstream analyses. Following this expression-based filtering step, differential expression analysis was performed using DESeq2 (v1.30.1) on the corresponding raw count matrix. Principal component analysis (PCA) was performed on variance-stabilized count data to assess sample clustering. Differentially expressed genes (DEGs) were defined as genes with an absolute log2 fold change ≥1 (corresponding to ≥2-fold change) and a false discovery rate (FDR) <0.01, following Benjamini-Hochberg correction.

### 2.8 Functional and pathway enrichment analysis

Gene Ontology (GO) Biological Process enrichment analysis of differentially expressed genes (DEGs) was performed using STRING v12.0, with the *Mus musculus* genome specified as the background. Acute and chronic DEG sets were analysed separately. GO terms were grouped by semantic similarity (≥0.7) to reduce redundancy, and statistical significance was assessed using FDR-adjusted *p*-values. Kyoto Encyclopedia of Genes and Genomes (KEGG) pathway enrichment analysis was performed independently for the acute and chronic DEG sets using the BMKCloud analysis platform (BMKGene). The proportion of DEGs assigned to individual KEGG pathways was used to compare pathway representation within each disease stage.

### 2.9 Extracellular matrix/fibrosis gene expression score

To assess coordinated extracellular matrix (ECM) and fibrosis-associated transcription, an a priori panel of 19 genes with established roles in ECM deposition, organization, cross-linking, remodelling, and fibrotic responses was examined: structural ECM/collagen genes (*Col1a1*, *Col1a2*, *Col3a1*, *Col4a1* and *Col6a1*), fibrosis- and myofibroblast-associated genes (*Acta2*, *Postn*, *Ctgf*, *Fn1* and *Tgfb1*), ECM cross-linking genes (*Loxl1*, *Loxl2* and *Loxl3*), matrix remodelling/regulatory genes (*Mmp2*, *Mmp9* and *Timp2*), and ECM organization-associated genes (*Lum*, *Dcn* and *Mfap5*).

RNA-seq expression values for each gene and individual mouse were transformed as log2(FPKM + 1). For calculation of the composite ECM fibrosis score, log2-transformed expression values were standardized separately for each gene across the complete cohort of 15 mice (uninfected, n = 5; chronically infected, n = 10). For each gene, the mean expression and sample standard deviation were calculated across all 15 mice, with the sample standard deviation determined using the STDEV.S function in Microsoft Excel. The standardized expression value for each individual mouse was then calculated as a Z-score according to Z=(x-\bar{x})/s, where x is the log2(FPKM + 1) expression value for an individual mouse, \bar{x} is the mean log2(FPKM + 1) expression of that gene across all 15 mice and is the corresponding sample standard deviation. Positive and negative Z-scores therefore represent expression above and below the cohort mean for the respective gene, respectively. A composite ECM fibrosis score was subsequently calculated for each mouse as the arithmetic mean of its Z-scores across all 19 genes.

Gene-level expression patterns were visualized using Heatmapper 2. The log2(FPKM + 1)-transformed expression matrix was uploaded to Heatmapper and scaled by row to visualize relative expression of each gene across individual samples. Hierarchical clustering of genes was performed using correlation distance and average linkage, and the resulting row dendrogram was displayed. Column clustering was disabled to preserve the predefined sample order, with uninfected mice (U1-U5) followed by chronically infected mice (I1-I10). Relative expression was displayed using a blue-white-red colour scale.

Composite ECM fibrosis scores were compared between uninfected and chronically infected mice using a two-tailed Mann-Whitney test in GraphPad Prism 11. Statistical significance was defined as P < 0.05.

### 2.10 Heatmap analysis

Heatmaps were generated using the web-based tool Heatmapper2. Input gene expression data were normalized by row using z-score transformation to facilitate comparison of relative expression patterns across samples. Hierarchical clustering was performed using correlation-based distance with average linkage. A custom blue-white-red colour scale with 50 bins was applied to visualize expression levels. Row dendrograms were displayed to represent clustering of features, whereas column dendrograms were not shown. No interpolation was applied. Unless otherwise specified, default parameters of Heatmapper2 were used.

### 2.11 Protein association network analysis

Protein association networks were constructed using STRING v12.0. Upregulated DEGs identified during acute (11 dpi) and chronic (171 dpi) infection were analysed separately, using *Mus musculus* as the reference organism. Networks were generated using the full STRING network and the highest-confidence minimum required interaction score (combined STRING score ≥0.9). All STRING evidence channels were enabled, including text mining, experiments, curated databases, co-expression, gene neighbourhood, gene fusion, and gene co-occurrence. No additional first- or second-shell interactors were added; therefore, networks contained only proteins encoded by the submitted DEG sets. Disconnected nodes were hidden from the network visualizations. Networks were displayed in confidence mode, in which edges represent STRING functional protein associations and edge thickness reflects the combined confidence score. The same network parameters were applied independently to the acute and chronic datasets.

## 3 Results

### 3.1 Anatomically standardised transcriptional profiling of the cardiac response to acute and chronic *Trypanosoma cruzi* infection

We used BLI imaging to longitudinally track *T. cruzi* dissemination and persistence (Lewis and Kelly, 2016; Lewis et al., 2016; Ward et al., 2020; Ward et al., 2020). In vivo imaging captures infection at a whole-animal level, in a non-invasive manner, while end-point ex vivo imaging provides information on tissue distribution and sites of parasite persistence (Figure 1). During the acute stage (Figure 1B and C), parasites were widely disseminated across multiple organs and tissues, including the heart (Figure 1C). At a cellular level, histological analysis confirmed the presence of intracellular *T. cruzi* amastigotes within cardiomyocytes (Figure 2A). As infection transitioned to the chronic stage, the parasite burden diminished substantially, with detectable BLI foci restricted predominantly to the skin and gastrointestinal tract (Figure 1D), in line with previous studies using the BALB/c:CL Brener infection model (Lewis et al., 2016; Ward et al., 2020; Lewis et al., 2014). In this model, chronic stage cardiac infection is intermittent, with an end-point detection frequency of approximately 10% (Lewis and Kelly 2016; Lewis et al., 2016; Lewis et al., 2014). We compared the cardiac transcriptional response in hearts during acute and chronic infection with those from uninfected mice. For an unbiased transcriptomic comparison, we implemented an anatomically standardized cardiac sampling strategy (Figure 2B and C). In the acute stage, BALB/c mice (n=10 infected, n=5 uninfected) were first examined by ex vivo imaging at 11 dpi, when all animals displayed extensive cardiac infection (Figure 1). A standardised 3 mm cardiac biopsy from the mid-right ventricle was then collected (Figure 2B and C) to minimise anatomical sampling variability between animals. At the chronic stage (171 dpi), although each of the mice remained infected, none displayed detectable cardiac bioluminescence at the point of analysis, under conditions in which the reported limit of detection is <20 parasites (Ward et al., 2020).

**Figure 1.**
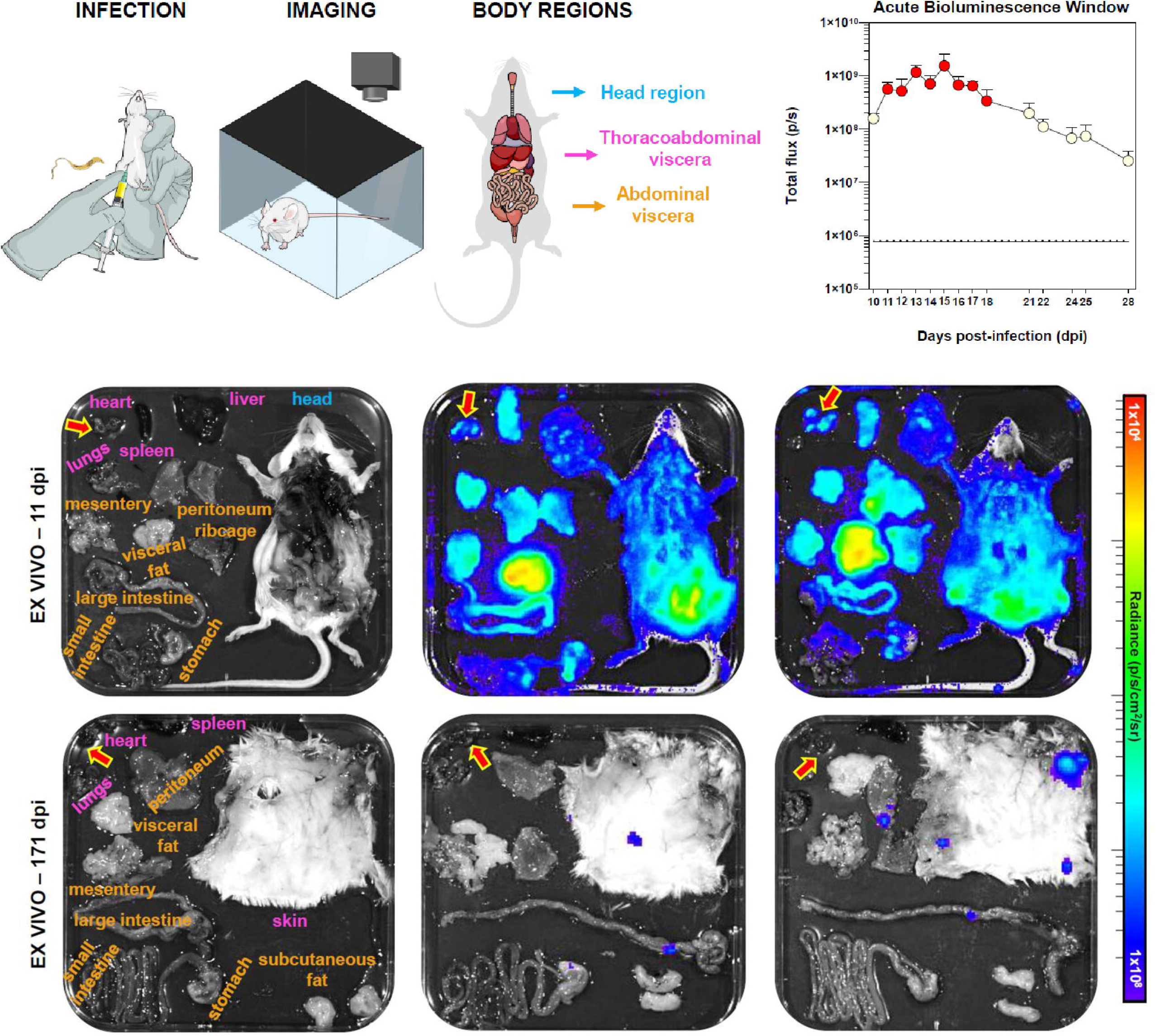
Parasite distribution during acute and chronic infection. **(A)** Schematic overview of workflow: BALB/c mice were infected i.p. with 1×10³ *T. cruzi* CL-Luc trypomastigotes and monitored by in vivo and ex vivo imaging (Material and Methods). **(B)** Longitudinal monitoring of whole-body bioluminescence during the acute stage of infection. Peak parasite burden (red dots) occurs 11-18 days post-infection (dpi). Data are mean ± SD. Horizontal black line indicates mean background bioluminescence from uninfected mice (n=3), dashed line indicates ± SD. **(C)** Ex vivo whole-body bioluminescence imaging at 11 dpi (n=10) showing widespread parasite distribution across organs and tissues, including the heart (indicated by arrow). **(D)** Ex vivo imaging of chronically infected mice at 171 dpi (n=10) showing absence of cardiac bioluminescence. Bioluminescence images are displayed as radiance (photons s ¹ cm ² sr ¹) using the colour scale shown. Imaging acquisition and display settings (exposure time, binning, and scale) were kept constant across all mice.

**Figure 2.**
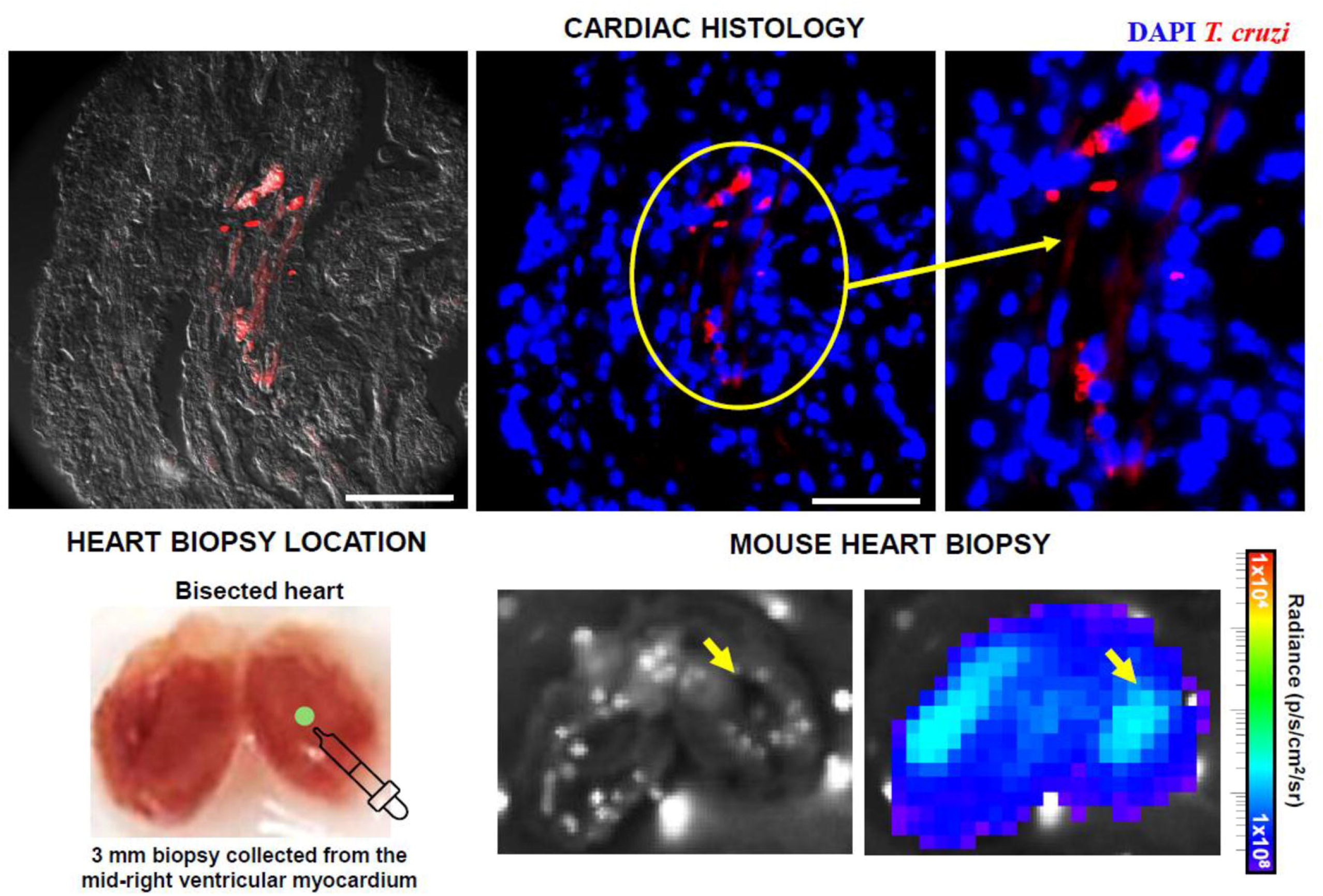
Imaging-guided anatomically standardized sampling of cardiac tissue from *T. cruzi* infected mice. **(A)** Acute stage cardiac infections at a cellular level. BALB/c mice were infected intraperitoneally with 1×10³ *T. cruzi* CL-Luc::mScarlet trypomastigotes. Representative cardiac histology sections from 14 dpi show intracellular *T. cruzi* amastigotes expressing mScarlet (magenta) within cardiomyocytes. Host nuclei are counterstained with DAPI (blue). Scale bars=50 μm. **(B)** Schematic illustrating the sampling of 3 mm mid-right ventricular myocardium biopsies collected following ex vivo imaging (Material and Methods). **(C)** Enlarged views of a bisected heart from a mouse infected with *T. cruzi* CL-Luc trypomastigotes (acute stage, 11 dpi; n=10) showing parasite distribution across cardiac tissue. Colour-matched arrows indicate the biopsy region used for RNA extraction. The bioluminescence image is displayed as radiance (photons s ¹ cm ² sr ¹) using the colour scale shown.

### 3.2 Comparison of the cardiac transcriptional profile during acute and chronic infection

RNA-seq data derived from the targeted cardiac biopsies (Figure 2B and C; Material and Methods) revealed disease stage-specific transcriptional changes during *T. cruzi* infection. PCA demonstrated clear segregation between infected and uninfected hearts (Figure 3; Supplementary Figure 1), consistent with pronounced infection-associated transcriptional remodelling at 11 dpi (Figure 3B). Separation between infected and uninfected samples was also observed at 171 dpi (Figure 3C), demonstrating persistent transcriptional differences during chronic infection. Differential expression analysis identified 64 upregulated and 9 downregulated genes during acute infection (log FC ≥1, FDR <0.01), that reflected broad activation of immune and inflammatory pathways (Figure 3D; Table 1; Supplementary Table 1; Supplementary Figure 2). At 171 dpi, 25 genes were upregulated and 2 were downregulated using the same statistical thresholds (Figure 3E; Table 1; Supplementary Table 1; Supplementary Figure 2), demonstrating that a significant cardiac transcriptional response persisted during chronic infection. Table 1 presents selected representative DEGs from the acute and chronic stages, highlighting genes associated with imune activation, antigen presentation and inflammatory responses. The complete DEG list for both stages are provided in Supplementary Table 1.

**Figure 3.**
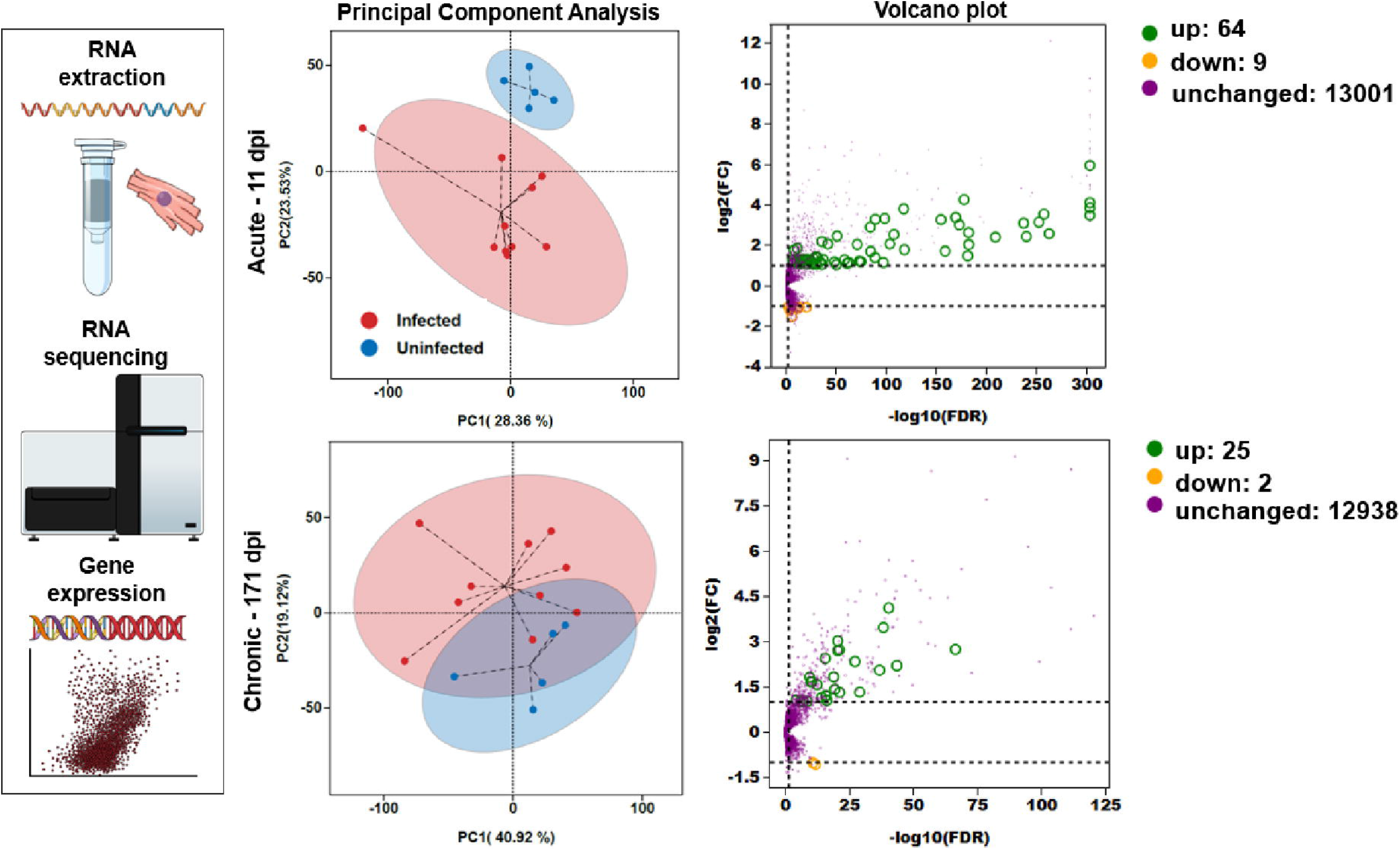
Stage-specific cardiac transcriptional responses during acute and chronic *T. cruzi* infection. **(A)** Overview of RNA extraction, sequencing, and downstream differential expression analysis from standardized cardiac biopsies. **(B-C)** Principal component analysis (PCA) of normalized gene expression profiles from infected (red, n=10) and uninfected control (blue, n=5) hearts at each stage. Acute infection (11 dpi) data show separation between infected and control samples. Chronic infection (171 dpi) data also segregate by infection status, suggestive of continued but reduced transcriptional divergence. Genes with expression values ≤5 FPKM were excluded prior to PCA and differential expression analysis. **(D-E)** Volcano plots of DEGs identified within each timepoint (log FC ≥1, FDR < 0.01). Each point represents one gene; the x-axis shows -log (FDR) and the y-axis log fold change. Dashed lines indicate statistical and fold-change thresholds. Purple points represent genes that did not meet the FDR cutoff despite apparent fold changes and therefore not classified as differentially expressed.

**Table 1.** Selected differentially expressed genes representing major transcriptional signatures in the murine heart during acute and chronic *T. cruzi* infection. Genes in bold are significantly upregulated at both the acute (11 dpi) and chronic (171 dpi) stages of infection and are designated as components of a conserved transcriptional signature. The directions (up/down) of differential gene expression are indicated by arrows. For more details see Supplementary Table 1.

| Antigen Presentation Signature | Interferon and Innate Immune Signature | Complement and Myeloid Activation | Oxidative Stress and Metabolic Adaptation |
|---|---|---|---|
| ***B2m*** **↑↑ Conserved** | ***Ifi27*** **↑↑ Conserved** | *C1qa* ↑− Acute-specific | ***Xdh*** **↑↑ Conserved** |
| ***Cd74*** **↑↑ Conserved** | ***Irf1*** **↑↑ Conserved** | *C1qb* ↑− Acute-specific | *Akr1b3* ↑− Acute-specific |
| ***H2-Aa*** **↑↑ Conserved** | ***Irgm1*** **↑↑ Conserved** | *C1qc* ↑− Acute-specific | *Pla2g16* ↑− Acute-specific |
| ***H2-D1*** **↑↑ Conserved** | ***Samhd1*** **↑↑ Conserved** | *C1ra* ↑− Acute-specific | *Nr1d1* ↓− Acute-specific |
| ***H2-K1*** **↑↑ Conserved** | ***Ube2l6*** **↑↑ Conserved** | *C1s1* ↑− Acute-specific | *Nudt13* ↑− Acute-specific |
| ***H2-Q4*** **↑↑ Conserved** | ***Wars*** **↑↑ Conserved** | *Ctss* ↑− Acute-specific | *Qdpr* ↓− Acute-specific |
| ***H2-T23*** **↑↑ Conserved** | ***Bst2*** **↑↑ Conserved** | *Tyrobp* ↑− Acute specific | *G0s2* ↑− Acute-specific |
| *H2-T22* −↑ Chronic-specific | ***Lgals3bp*** **↑↑ Conserved** | *Fgl2* ↑− Acute-specific | *Rrad* ↑− Acute-specific |
| *Tapbp* ↑− Acute-specific | *Ifi27l2a* ↑− Acute-specific | ***Ly6a*** **↑↑ Conserved** | Extracellular Matrix and Tissue Remodelling |
| Immunoproteasome and Antigen Processing | *Ifi35* ↑− Acute-specific | *Ly6c1* ↑− Acute-specific | *Col3a1* ↓− Acute-specific |
| ***Psmb10*** **↑↑ Conserved** | *Ifitm3* ↑− Acute-specific | *Ly6c2* ↑− Acute-specific | *Col4a1* ↓− Acute-specific |
| ***Psme2*** **↑↑ Conserved** | *Irf9* ↑− Acute-specific | *Ly6e* ↑− Acute-specific | *Lum* ↓− Acute-specific |
| ***Psme2b*** **↑↑ Conserved** | *Stat3* ↑− Acute-specific | *C3* −↑ Chronic-specific | *Ltbp4* ↓− Acute-specific |
| *Pomp* ↑− Acute-specific | *Trafd1* ↑− Acute-specific | **Legend:** | *Vwa5a* ↑− Acute-specific |
| *Psme1* ↑− Acute-specific | *Uba7* ↑− Acute-specific | **↑↑ = conserved upregulated** | *Mtap5* ↓− Acute-specific |
| Chronic Cardiac Pathology Markers | *Sp100* ↑− Acute-specific | **↑ – = acute only** |  |
| *Nppb* −↓ Chronic-specific | *Stat1* −↑ Chronic-specific | **– ↑ = chronic only** |  |
| *Tnfrsf12a* −↓ Chronic-specific |  |  |  |

**Footnote: acute and chronic datasets were analysed independently relative to their corresponding uninfected controls; fold-change estimates should therefore be interpreted within each disease stage rather than as direct statistical comparisons between stages.**

Comparison of the acute and chronic DEG sets revealed a subset of 20 transcripts that was consistently upregulated at both stages, defining a conserved transcriptional profile dominated by genes associated with antigen presentation, IFN signalling, innate immune activation, and immunoproteasome function (Table 1). Within the antigen presentation group, *B2m*, *Cd74*, *H2-Aa*, *H2-D1*, *H2-K1*, *H2-Q4*, and *H2-T23* were significantly upregulated at both stages, indicating sustained activation of MHC-dependent immune pathways throughout disease progression. Similarly, the immunoproteasome-associated genes *Psmb10*, *Psme2*, and *Psme2b* remained upregulated, suggesting sustained enhancement of antigen-processing capacity within cardiac tissue. The IFN-inducible genes and genes involved in innate immune and antiviral responses that were conserved across both disease states (Cheung et al., 2025; Xu et al., 2019) included *Ifi27*, *Irf1*, *Irgm1*, *Samhd1*, *Ube2l6*, *Wars*, *Bst2*, and *Lgals3bp* (Table 1).

Biological Process Gene Ontology (GO) enrichment analysis revealed distinct patterns of function enrichment during acute infection (Figure 4). As described above, DEGs were broadly enriched for antigen processing and presentation pathways, including both MHC class I and II processes, alongside T cell-mediated cytotoxic activity, IFN responses, and a range of innate and adaptive immune pathways (Figure 4A). In contrast, the chronic stage DEG set was more restricted (Figure 4B). Heatmap analysis further resolved these transcriptional patterns (Figure 5). In the acute infection (Figure 5A), immune- and IFN-associated genes showed heterogeneous expression patterns among individual mice. During the chronic stage (Figure 5B), a more restricted set of immune- and IFN-associated genes remained upregulated. Collectively, GO (Figure 4), KEGG (Supplementary Figure 3) analyses, together and gene-level heatmap visualization (Figure 5), indicate that the chronic cardiac transcriptional response is reduced compared to the acute response.

**Figure 4.**
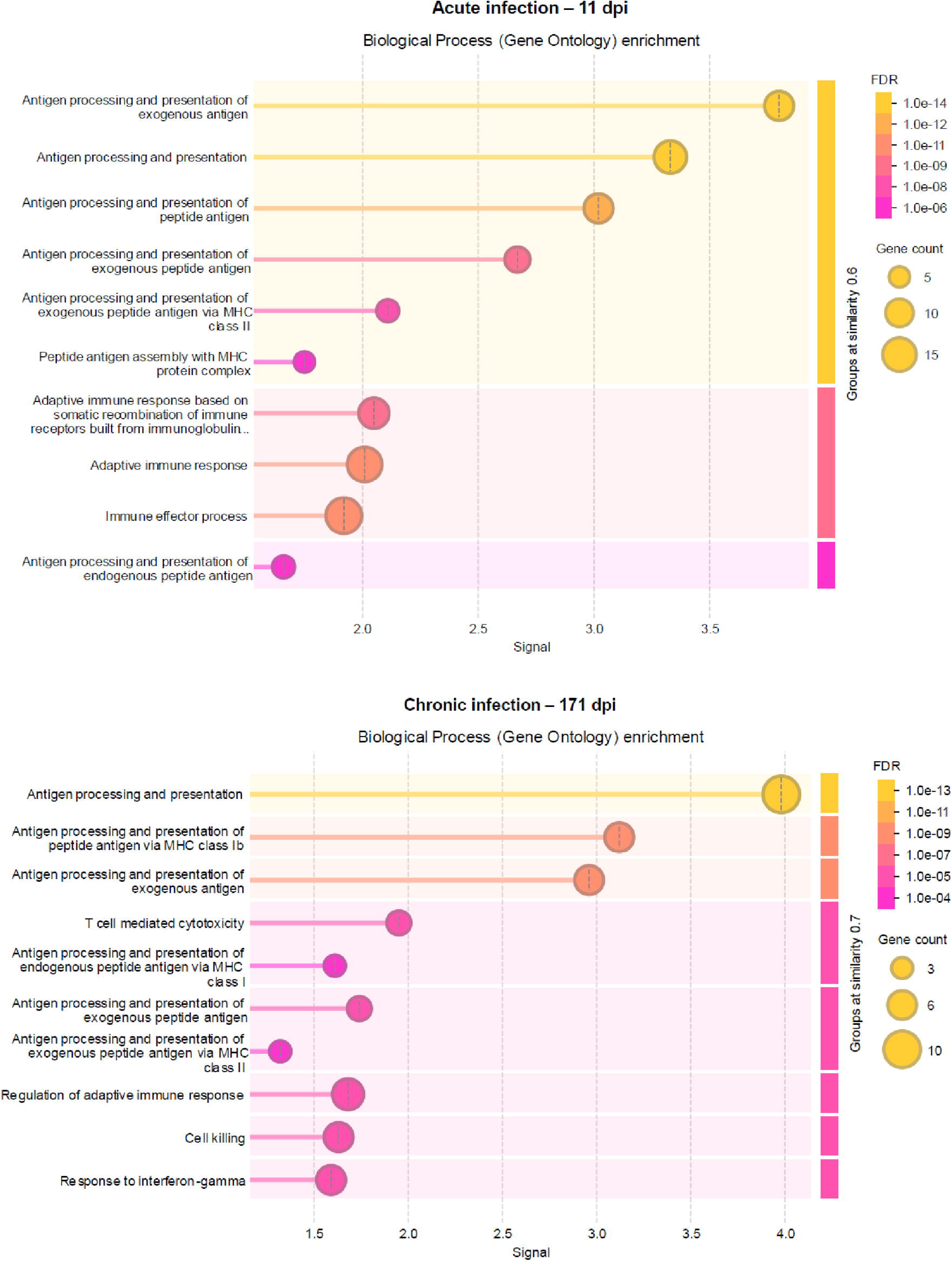
Gene Ontology (GO) Biological Process enrichment reveals a shift from broad immune activation in acute infection to a more focused profile during the chronic stage. **(A)** Acute stage infection (11 dpi) and **(B)** chronic stage infection (171 dpi) DEGs (log FC ≥1, FDR <0.01) were analysed for GO Biological Process enrichment using STRING (v12.0), with the mouse genome as background. GO terms were grouped by semantic similarity (≥0.7) to reduce redundancy. The x-axis represents enrichment signal (-log10 FDR-adjusted p-value), where higher values indicate stronger statistical overrepresentation. Bubble size indicates the number of DEGs associated with each GO term, and colour denotes the FDR.

**Figure 5.**
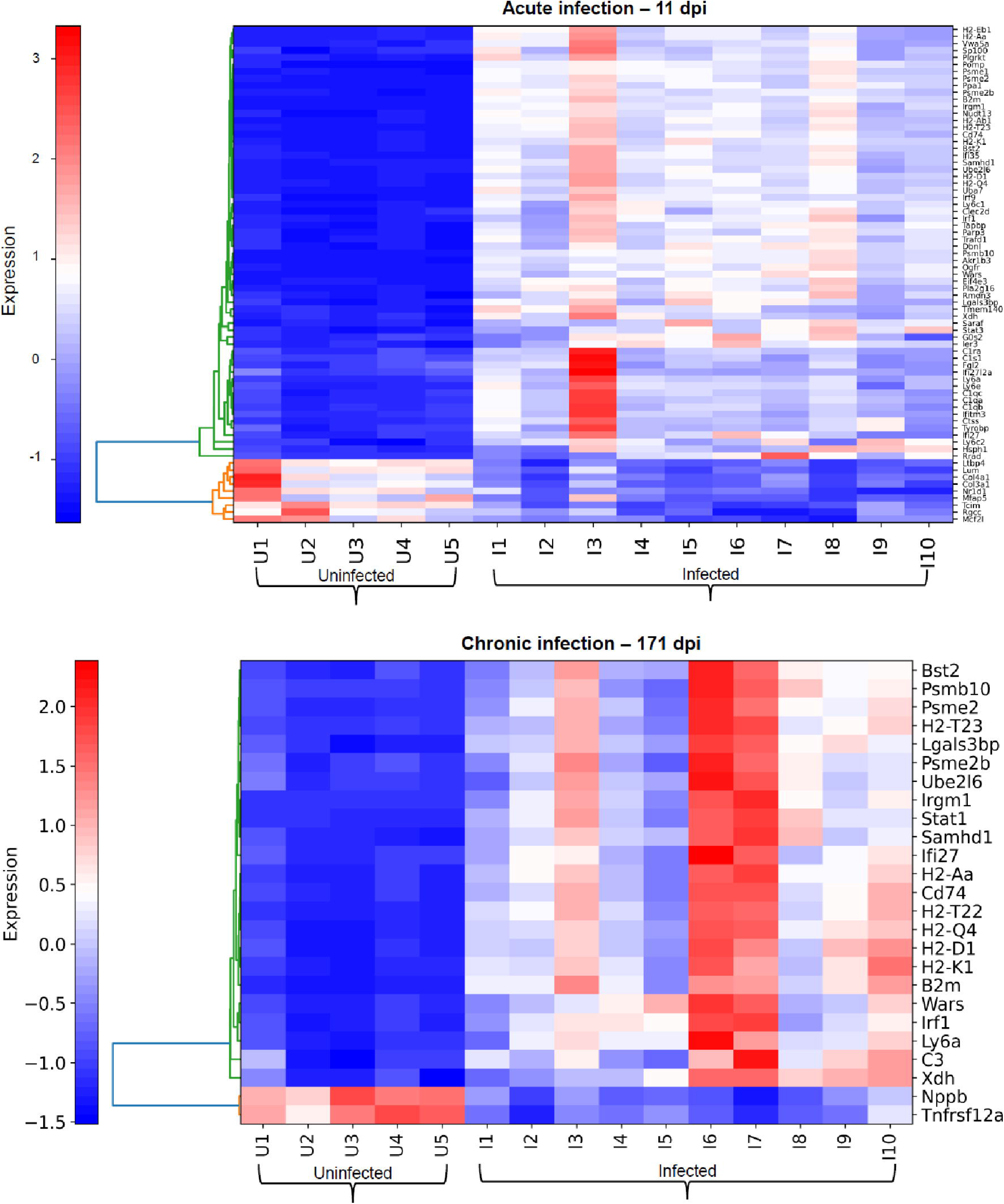
Stage-specific transcriptional signatures distinguish infected and uninfected mouse hearts. Heatmaps of DEGs (log FC ≥1, FDR <0.01) at each stage of infection. **(A)** Acute infection (11 dpi): gene expression profiles across individual uninfected (U1-U5) and infected (I1-I10) mice. **(B)** Chronic infection (171 dpi): DEG profiles at the chronic stage across uninfected (U1-U5) and infected (I1-I10) mice. Rows represent genes and columns represent individual samples. Expression values are displayed as row scaled Z-scores, enabling comparison of relative expression patterns. Uninfected and infected samples cluster separately. Complete DEG lists are provided in Table S1. Heatmap generated using Heatmapper2 (row z-score normalization; correlation distance; average linkage).

Several IFN-stimulated genes, including *Ifi27l2a*, *Ifi35*, *Ifitm3*, *Irf9*, *Trafd1*, *Uba7*, *Sp100* and *Stat1*, were induced exclusively at 11 dpi, indicating a broader IFN-associated transcriptional response during the acute stage. This time point was also characterised by enhanced expression of genes associated with metabolic regulation and cellular stress responses (Table 1, Supplementary Table 1), including those involved in lipid metabolism (*G0s2* and *Pla2g16*), oxidative stress (*Akr1b3*), apoptosis and metabolic regulation (*Nudt13*, *Stat3* and *Hsph1*), translational control (*Eif4e3*), and calcium/endoplasmic reticulum stress responses (*Rrad* and *Saraf*). Genes associated with complement activation and inflammatory myeloid-cell recruitment showed a similar temporal pattern, including members of the classical complement pathway (*C1qa*, *C1qb*, *C1qc*, *C1ra*, and *C1s1*) together with genes associated with monocyte and macrophage activation (*Ctss*, *Tyrobp*, *Fgl2*, *Ly6c1*, *Ly6c2* and *Ly6e*). Among these genes, *Ly6a* remained significantly upregulated in both disease stages, indicating persistence into the chronic stage. Collectively, these findings highlight prominent complement- and myeloid-associated transcription during acute infection (Table 1, Supplementary Table 1).

To examine functional associations among proteins encoded by the differentially expressed genes, STRING analysis was performed using all significantly upregulated DEGs identified at each infection stage (Table 1). Highest-confidence protein association networks (combined score ≥0.9) revealed distinct stage-associated organisation of the cardiac response (Figure 6).

**Figure 6.**
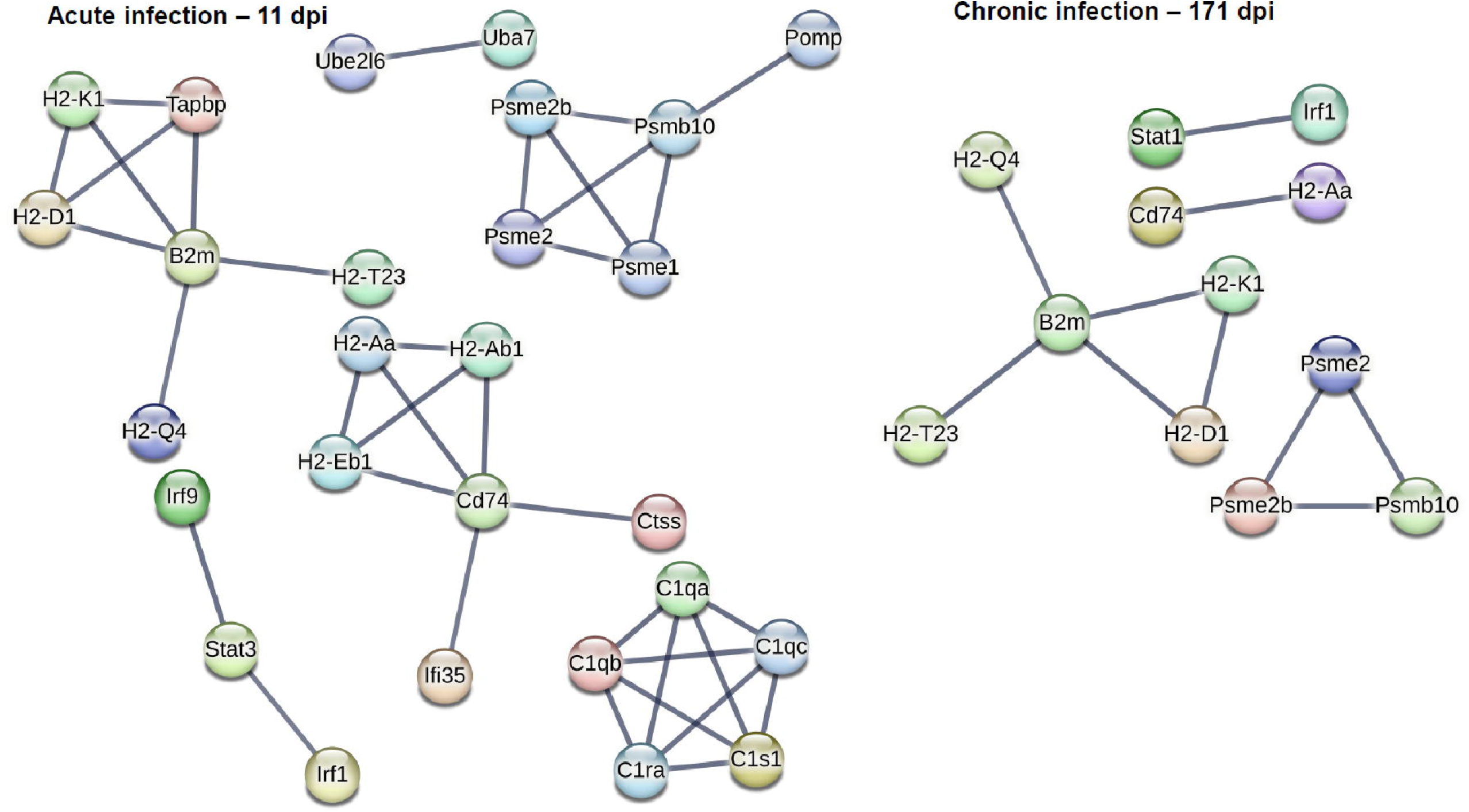
Protein association networks during acute and chronic *T. cruzi* infection. STRING networks (v12.0) were generated separately from significantly upregulated differentially expressed genes (DEGs; log2FC ≥ 1, FDR < 0.01) identified at each infection stage,, rather than from the representative subset presented in Table 1. Associations were restricted to the STRING highest-confidence threshold (combined score ≥ 0.9), with all evidence channels enabled and no additional interactors added. Networks are shown for **(A)** acute infection (11 dpi) and **(B)** chronic infection (171 dpi), using identical network parameters. Nodes represent proteins encoded by the submitted upregulated DEGs, and edges represent STRING functional protein associations meeting the specified confidence threshold; edge thickness reflects the combined confidence score. Disconnected nodes were hidden from visualization and therefore submitted proteins without an association to another query protein at the specified threshold are not displayed.

The acute network (Figure 6A) contained several interconnected immune-associated clusters. Antigen processing and presentation included MHC class I and II components such as *H2-K1*, *H2-D1*, *H2-Aa*, *H2-Ab1*, *H2-Eb1*, *H2-Q4* and *H2-T23*, together with *B2m*, *Cd74* and *Tapbp*. A distinct immunoproteasome-associated module included *Psmb10*, *Psme1*, *Psme2* and *Psme2b*, while a highly interconnected complement module comprised *C1qa*, *C1qb*, *C1qc*, *C1ra* and *C1s1*. IFN-associated proteins, including *Irf1*, *Irf9*, *Ifi35* and *Stat3*, were also represented within the acute network. Additional connected proteins are shown in Fig 6, with the complete DEG dataset provided in Supplementary Table 1.

In contrast, the chronic stage network (Figure 6B) contained fewer connected proteins at the same STRING confidence threshold but retained a prominent antigen processing and presentation component centred on *B2m*, together with *Cd74* and MHC-associated proteins including *H2-K1, H2-D1, H2-Aa, H2-Q4* and *H2-T23*. An immunoproteasome-associated module comprising *Psme2, Psme2b* and *Psmb10* was also retained, together with the IFN-associated proteins *Stat1* and *Irf1*. Notably, the complement-associated module evident during acute infection was absent in the chronic network. Thus, although the composition of the STRING network differed between stages, antigen processing and presentation and immunoproteasome-associated proteins remained prominent components of the transcriptional response during chronic infection.

### 3.3 Limited transcriptional evidence for widespread cardiac remodelling during acute and chronic infection

Downregulation of several ECM-associated transcripts at 11 dpi, including *Col4a1*, *Col3a1*, *Lum*, and *Mtap5* (Figure 3D; Table 1; Supplementary Table 1), indicated that classical ECM-remodelling programmes were not broadly activated during the acute stage of infection. Downregulation of *Rgcc* and *Tcim* (Supplementary Table 1) further supported altered regulation of pathways associated with stromal, inflammatory, and tissue-remodelling responses during acute infection.

At the chronic stage, the transcriptional profile of the anatomically defined cardiac region examined in this study similarly provided limited evidence for widespread ECM/fibrosis-associated activation. *Tnfsf12a* (Fn14; fibroblast growth factor-inducible protein 14) and *Nppb* (the B-type natriuretic peptide), genes associated with inflammatory signalling and cardiac stress responses, respectively, were among the principal downregulated transcripts at 171 dpi (Figure 3E; Supplementary Table 1). To further assess coordinated ECM-associated transcription, expression of a 19-gene ECM/fibrosis-associated panel was examined across individual animals. The composite ECM fibrosis score did not differ significantly between uninfected and chronically infected mice (Mann-Whitney U = 19, two-tailed P = 0.514; Supplementary Figure 4). Similarly, the corresponding heatmap showed heterogeneous expression of individual ECM/fibrosis-associated genes across mice, without a uniform coordinated increase across chronically infected samples.

## 4 Discussion

The development of Chagas cardiomyopathy is a prolonged and heterogeneous process in which parasite persistence and the host response interact over many years. Experimental models have been central to resolving these relationships because they permit parasite burden, tissue distribution and pathology to be followed in ways that are not possible in humans (Lewis and Kelly, 2016; Lewis et al., 2016). Current evidence strongly suggests that parasite persistence is required for disease progression, with earlier hypotheses postulating a central role for autoimmunity no longer widely supported (Lewis et al., 2016; Macaluso et al., 2023; Puerta et al., 2023). Similarities between Chagasic cardiac pathology in humans and mice have led to experimental murine models becoming a key component of research on disease pathogenesis (Khan et al., 2024). In mice, continuous re-infection of the heart by parasites trafficked from immunotolerant sites, such as the GI tract, has been proposed to induce localised immune responses that eliminate infection foci, but result in cumulative collateral damage to surrounding tissue (Lewis and Kelly, 2016). The low regenerative capacity of cardiac tissue (Francisco et al., 2018) makes the heart particularly vulnerable to this type of progressive pathology. It should be noted that cardiac disease in humans does not completely overlap with that in mice. Arrhythmias are a hallmark of human *T. cruzi* infection and are frequently associated with progressive cardiac dysfunction leading to heart failure (Nunes et al., 2023; Nunes et al., 2024). In contrast, the severity of *T. cruzi* induced arrhythmias varies considerably between mouse strains. With BALB/c mice, the impact of infection on cardiac function is limited largely to bradycardia in the acute stage, although the development of fibrosis during chronic infections is a common outcome (Francisco et al., 2018; Francisco et al., 2023).

Against this background, the principal finding of the present study is that the marked reduction in detectable cardiac parasite burden is accompanied by contraction, but not complete resolution, of the cardiac transcriptional response. A 20-gene set remained significantly upregulated in tissue sampled from a defined mid-right ventricle location.

This signature was upregulated at both the acute and chronic time points, despite the absence of detectable cardiac bioluminescence in the animals sampled at the chronic stage. Although activation of immune pathways during acute *T. cruzi* infection has been well established (Gironès et al., 2014; Ferreira et al., 2014; Venturini et al., 2023), the continued upregulation of a coordinated set of genes that are involved in antigen presentation, IFN signalling and immunoproteasome activity, throughout disease progression, has not previously been described. The most prominently conserved signature involved MHC-related genes together with immunoproteasome components. Persistent transcription could therefore reflect episodic cardiac exposure to parasites or parasite-derived antigens, transient reseeding below the detection window, systemic immune stimulation arising from parasites persisting elsewhere, or a locally maintained immune state following earlier infection (Bonney and Engman, 2008). These alternatives are not mutually exclusive. Irrespective of the underlaying mechanism, the data show that a restricted immune-associated programme remains detectable in cardiac tissue even when the local parasite burden is extremely low or intermittent (Pérez-Mazliah et al., 2021). A second key feature of the conserved signature was the maintenance of an IFN-associated transcriptional network, involving both type I and II interferons, comprising *Ifi27*, *Irf1*, *Irgm1*, *Samhd1*, *Ube2l6*, Bst2, *Lgals3bp*, and *Wars* (Xu et al., 2019; Zhao et al., 2025; Goswami et al., 2022; Murray et al., 2015; Wang et al., 2026; Wang et al., 2026). IFN signalling plays an essential role in controlling intracellular *T. cruzi* replication through activation of macrophage effector mechanisms and promotion of Th1 immunity. However, prolonged induction of IFN activity may also contribute to tissue damage through sustained inflammatory signalling (Ferreira et al., 2014; Wen et al., 2025).

Upregulation of genes such the antiviral restriction factor *Samhd1* (Cheung et al., 2025), which acts to reduce the intracellular dNTP pool, could be a general adaptative response to infection, while increased expression of the galectin binding protein gene *Lgals3bp* (Xu et al., 2019) may be linked with modulation of immune cell interactions within cardiac tissue. The expression profile of *Irgm1,* which was amongst the most strongly upregulated genes (Table 1), highlighted a potential role for IFN-inducible GTPases in cardiac host defence during both acute and chronic infection. While *Irgm1* has been implicated in resistance to intracellular pathogens in other infection models (Goswami et al., 2022), its induction in cardiac tissue during *T. cruzi* infection suggests a potentially underappreciated effect resulting from host-pathogen interaction. *Irgm1* also has a broader role in regulating immune cell function. Neutrophil specific deletion of the gene exacerbates cardiac dysfunction, impairs post-myocardial infarction repair, and disrupts neutrophil clearance through defective efferocytosis (Wang et al., 2026; Wang et al., 2026).

In contrast to the conserved immune programme, upregulation of genes associated with complement activation and inflammatory myeloid recruitment was largely restricted to acute infection, the stage most associated with myocarditis (Gironès et al., 2014), and with bradycardia in this mouse model (Francisco et al., 2023). Elevated expression of C1q family members, *Ctss*, *Tyrobp*, *Fgl2*, *Ly6c1*, and *Ly6c2* could reflect the intense innate immune response required to control acute stage infections, during which intracellular parasite replication is more rapid than during the chronic stage (Ward et al., 2020). The subsequent decline in expression of these genes during chronic infection suggests a transition from broad innate inflammatory activation toward a more focused adaptive immune response and is consistent with the reduced inflammatory state of cardiac tissue observed during the chronic stage (Cunha-Neto et al., 2005; Nunes et al., 2021; Laugier et al., 2020). However, inferences about the whole-heart response should remain tentative at this stage, given the localised nature of the tissue biopsies that were sampled, and the temporal impact of infection dynamics (Lewis et al., 2026).

Distinct disease stage-specific regulation was also observed in pathways associated with oxidative stress and metabolic adaptation, with xanthine dehydrogenase (XDH) being the only DEG in this functional group to exhibit conserved upregulation throughout infection. XDH plays a central role in purine oxidative metabolism but can also contribute to the production of reactive oxygen species and nitric oxide following conversion of the enzyme to its alternative catalytic form, xanthine oxidase (XOR) (Nelson and Voruganti, 2026). Heart tissue in *T. cruzi* infected mice is characterised by depletion of purine nucleotides and accumulation of the downstream metabolic products xanthosine, xanthine and urate (Polito et al., 2021). This may impact the redox status of infected tissue since increased levels of xanthine and urate give rise to the generation of hydrogen peroxide. Enhanced expression of XOR has also been implicated in inflammatory heart disease through perturbation of the cardiac redox balance (Gironès, et al., 2014). The antiviral restriction factor SAMHD1, which is also upregulated in both stages of the disease, may act in concert with XDH/XOR, since it acts to deplete dNTP levels (Ferreira et al., 2014), although its expression is not associated with enhanced oxidative stress.

A notable observation from our data was that upregulation of the transcription factor *Stat1,* a crucial effector of cellular responses to both type I and II interferons (Wen et al., 2025), was specific to the chronic stage of infection. In contrast, upregulation of *Stat3*, a key mediator of the IL-10 associated pathways that moderate inflammatory immune responses, was confined to the acute stage (Sarma et al., 2021). This expression profile suggests a role for *Stat3* in reducing cardiac inflammation at the period when parasite levels are at their highest and the risk of myocarditis is greatest (Figures 1C and 2A), whereas chronic *Stat1* upregulation is consistent with continued Th1/IFN-g associated anti-parasitic activity in response to episodic *T. cruzi* exposure, which may also contribute to tissue pathology.

At the network level, these transcriptional patterns were reinforced by highest-confidence protein association networks that align with immune processes previously implicated in Chagas cardiomyopathy (de Castro et al., 2020; Navarro et al., 2015; Ferreira et al., 2017; Cunha-Neto et al., 2005; Nunes et al., 2021), including IFN-associated responses and antigen processing and presentation. However, previous evidence has largely relied on in vitro cardiomyocyte models or indirect datasets, such as microRNA studies (Navarro et al., 2015; Ferriera et al., 2017; Laugier et al., 2020), with comparatively limited characterisation of transcriptional responses within anatomically defined cardiac regions across both stages of infection.

Given the established roles of *Col4a1* in basement membrane organization and *Lum* in collagen fibrillogenesis and ECM assembly (Van Agtmael et al., 2005; Rixon et al., 2023), these changes suggest early disruption of ECM homeostasis rather than a uniform profibrotic transcriptional response. The lack of a coordinated ECM/fibrosis-associated transcriptional signature within this defined region is compatible with a model in which pathological remodelling is spatially heterogeneous, with focal areas of fibrosis or tissue injury potentially coexisting with regions showing comparatively limited transcriptional evidence of ECM activation. Several limitations should be considered. First, RNA-seq cannot distinguish altered transcription within resident cardiac cells from changes in the abundance or activation state of infiltrating immune populations. Single-cell or spatial transcriptomic approaches could help resolve the cellular sources of the conserved chronic programme. Second, the fixed anatomical biopsy improves reproducibility but necessarily samples only a small fraction of a spatially heterogeneous organ and may miss focal parasite-infected inflammatory lesions or fibrotic areas. Third, cardiac bioluminescence provides sensitive detection of parasite burden but captures only a temporal snapshot at the time of tissue collection.

Collectively, these findings support a model in which the cardiac response to *T. cruzi* infection evolves from widespread parasite-associated inflammation during the acute stage to a more restricted but persistent immune-associated transcriptional state during chronic infection. The latter is characterised by sustained antigen-presentation, IFN-associated and immunoproteasome transcription despite very low and spatially intermittent cardiac parasite burden. At the same time, the anatomically defined region examined here showed little evidence of coordinated active fibrotic transcription, emphasising that immune activation and ECM remodelling need not be spatially or temporally coincident. Thus, parasite containment during chronic infection can coexist with persistent cardiac immune activation, providing a molecular context for cumulative and spatially heterogeneous tissue injury over time.

## 6 Author Contributions

Conceptualization: AFF and JMK.

Data curation: AFF.

Formal analysis: AFF.

Funding acquisition: AFF, FO and JMK.

Investigation: AFF, FO and SJ.

Methodology: AFF and FO.

Project administration: JMK.

Resources: AFF, FO and SJ.

Supervision: AFF and JMK.

Validation: AFF.

Visualization: AFF, FO and SJ.

Writing - original draft: AFF and JMK.

Writing - review & editing: AFF, JMK, FO, MCT and SJ.

## 8 Acknowledgments

We would like to thank the Biological Service Facility team and Abdi Elmi at LSHTM for advice. Figures 1a and 3a were created using graphical elements from Science Figures (Open Design License 1.0) and the NIH BioArt and SciDraw galleries. RNA-seq data processing and analysis were performed using the BMKCloud platform provided by BMKGene (Beijing, China).

## 12 Data Availability Statement

Raw RNA-seq data supporting the findings of this study have been deposited in the European Nucleotide Archive (ENA) at EMBL-EBI under accession number PRJEB101446 (https://www.ebi.ac.uk/ena/browser/view/PRJEB101446) and are available. All processed datasets and source data underlying the figures presented in this study will also be made publicly available upon publication.

---

**Supplementary Figure 1. Global distribution and variability of transcriptome expression across acute and chronic cardiac samples. (A)** Density plots of log10-transformed FPKM values illustrating overall gene expression distributions for individual mouse heart samples collected at 11 dpi (acute stage; uninfected N1-N5, infected I1-I10) and 171 dpi (chronic stage; uninfected U1-U5, infected I1-I10). Overlapping density curves indicate comparable global transcript abundance distributions across samples within each stage. **(B)** Boxplots of log10(FPKM) values showing gene expression dispersion for each sample at acute (top) and chronic (bottom) stages. Similar median expression levels and interquartile ranges across samples confirm preserved library complexity and absence of major outliers, supporting data quality and cross-group comparability.

**Supplementary Figure 2. Pairwise sample-level differential expression and identification of recurrent transcriptional signatures. (A)** Acute stage infection (11 dpi): flower plot illustrating genes identified in differential expression analyses where individual uninfected mice (N1-N5) were each contrasted against the infected cohort (I1-I10). Each petal represents the number of DEGs detected in a specific single sample versus group comparison, and the central core indicates genes shared across comparisons. Bar plots (right) display total, upregulated, and downregulated genes identified per comparison. **(B)** Chronic stage infection (171 dpi): equivalent analyses in which individual uninfected mice (U1-U5) were contrasted against the infected cohort at the chronic stage, with shared genes shown in the central core and DEG counts per comparison summarized in bar plots. Importantly, DEG counts shown here derive from single sample versus group contrasts and therefore differ from the final group level differential expression results presented in Fig 3 and S1 Table, which were generated using statistical modelling across all biological replicates simultaneously. This analysis illustrates the reproducibility and consistency of infection-associated transcriptional signatures across individual animals.

**Supplementary Figure 3. Kyoto Encyclopedia of Genes and Genomes pathway enrichment analysis reveals sustained antigen presentation signatures during chronic cardiac infection.** Bar plots show Kyoto Encyclopedia of Genes and Genomes pathway enrichment of differentially expressed genes (DEGs) identified at each stage of infection. **(A)** Acute infection – 11 dpi: enriched pathways span infection and immunity-associated categories, including phagosome formation, cytokine pathway, JAK-STAT, complement cascades, and antigen processing and presentation, aligned with broad IFN driven innate immune activation and inflammation. **(B)** Chronic infection – 171 dpi: although fewer DEGs are detected overall, immune associated pathways remain enriched. Antigen processing and presentation pathways, including MHC class I and class II related signatures, represent a proportionally larger component of the chronic transcriptional response, indicating sustained engagement of antigen presentation and adaptive immune mechanisms. Percentages indicate the proportion of DEGs assigned to each Kyoto Encyclopedia of Genes and Genomes pathway within each stage.

**Supplementary Figure 4. Extracellular Matrix/fibrosis-associated transcriptional profile in hearts from individual chronically infected BALB/c mice at 171 days post-infection. (A)** Composite ECM (Extracellular Matrix) fibrosis score in uninfected (n = 5) and chronically infected (n = 10) mice. Gene expression values were log2-transformed and standardized to Z-scores for each gene, and the composite ECM fibrosis score for each mouse was calculated as the mean Z-score across the 19 ECM/fibrosis-associated genes. Individual points represent individual mice, and horizontal lines indicate group medians. No significant difference in ECM fibrosis score was detected between uninfected and chronically infected mice (two-tailed Mann-Whitney test, U = 19, P = 0.514). **(B)** Heatmap showing standardized expression (Z-scores) of the 19 ECM/fibrosis-associated genes across individual uninfected (U1-U5) and chronically infected (I1-I10) mice. Red indicates relatively higher expression and blue indicates relatively lower expression for each gene across samples.

**Supplementary Table 1.** Differentially expressed cardiac genes during acute and chronic *Trypanosoma cruzi* infection. The direction of differential gene expression (|log FC| ≥ 1, FDR < 0.01) compared to uninfected mice during acute (11 dpi) and chronic (171 dpi) stages of infection is shown. Biological description is summarised using standardised functional annotations.

| **Gene name** | **Acute<br>(11 dpi)** | **Chronic<br>(171 dpi)** | **Stage specificity** | **Biological description** |
|---|---|---|---|---|
| ***B2m*** | ↑ | ↑ | Acute+chronic | MHC I component |
| ***Bst2*** | ↑ | ↑ | Acute+chronic | Antiviral immune response |
| ***Cd74*** | ↑ | ↑ | Acute+chronic | MHC II processing |
| ***H2-Aa*** | ↑ | ↑ | Acute+chronic | MHC II / antigen presentation |
| ***H2-D1*** | ↑ | ↑ | Acute+chronic | MHC I / antigen presentation |
| ***H2-K1*** | ↑ | ↑ | Acute+chronic | MHC I / antigen presentation |
| ***H2-Q4*** | ↑ | ↑ | Acute+chronic | MHC I / antigen presentation |
| ***H2-T23*** | ↑ | ↑ | Acute+chronic | MHC I / antigen presentation |
| ***Ifi27*** | ↑ | ↑ | Acute+chronic | IFN-a response |
| ***Irf1*** | ↑ | ↑ | Acute+chronic | IFN-g signalling regulator |
| ***Irgm1*** | ↑ | ↑ | Acute+chronic | Immune defense / intracellular pathogen response |
| ***Lgals3bp*** | ↑ | ↑ | Acute+chronic | Immune modulation / galectin pathway / Type I/II IFN-response |
| ***Ly6a*** | ↑ | ↑ | Acute+chronic | Immune cell marker |
| ***Psmb10*** | ↑ | ↑ | Acute+chronic | Immunoproteasome / antigen processing |
| ***Psme2*** | ↑ | ↑ | Acute+chronic | Proteasome activation |
| ***Psme2b*** | ↑ | ↑ | Acute+chronic | Proteasome activation |
| ***Samhd1*** | ↑ | ↑ | Acute+chronic | Antiviral defence / nucleotide metabolism |
| ***Ube2l6*** | ↑ | ↑ | Acute+chronic | IFN-a/b induced ubiquitination |
| ***Wars*** | ↑ | ↑ | Acute+chronic | Protein synthesis / immune response |
| ***Xdh*** | ↑ | ↑ | Acute+chronic | Oxidative stress / metabolism |
| ***Akr1b3*** | ↑ |  | Acute | Oxidative stress / metabolism |
| ***C1qa*** | ↑ |  | Acute | Complement activation |
| ***C1qb*** | ↑ |  | Acute | Complement activation |
| ***C1qc*** | ↑ |  | Acute | Complement activation |
| ***C1ra*** | ↑ |  | Acute | Complement activation |
| ***C1s1*** | ↑ |  | Acute | Complement activation |
| ***C3*** |  | ↑ | Chronic | Complement activation |
| ***Clec2d*** | ↑ |  | Acute | Innate immune receptor |
| ***Col3a1*** | ↓ |  | Acute | ECM / fibrosis |
| ***Col4a1*** | ↓ |  | Acute | ECM / fibrosis |
| ***Ctss*** | ↑ |  | Acute | Antigen processing / lysosomal |
| ***Dbnl*** | ↑ |  | Acute | Actin cytoskeleton / cell structure |
| ***Eif4e3*** | ↑ |  | Acute | Protein translation |
| ***Fgl2*** | ↑ |  | Acute | Immune coagulation / inflammation |
| ***G0s2*** | ↑ |  | Acute | Lipid metabolism / cell regulation |
| ***H2-Ab1*** | ↑ |  | Acute | MHC II / antigen presentation |
| ***H2-Eb1*** | ↑ |  | Acute | MHC II / antigen presentation |
| ***H2-T22*** |  | ↑ | Chronic | MHC I / antigen presentation |
| ***Hsph1*** | ↑ |  | Acute | Stress response / heat shock |
| ***Ier3*** | ↑ |  | Acute | Stress response / immune activation |
| ***Ifi27l2a*** | ↑ |  | Acute | IFN-a/b response |
| ***Ifi35*** | ↑ |  | Acute | Type I/II IFN response |
| ***Ifitm3*** | ↑ |  | Acute | Type I/II IFN response |
| ***Irf9*** | ↑ |  | Acute | Type I/II IFN response |
| ***Ltbp4*** | ↓ |  | Acute | TGF-β regulation |
| ***Lum*** | ↓ |  | Acute | ECM |
| ***Ly6c1*** | ↑ |  | Acute | Immune cell marker |
| ***Ly6c2*** | ↑ |  | Acute | Immune cell marker |
| ***Ly6e*** | ↑ |  | Acute | Immune cell marker |
| ***Mcf2l*** | ↓ |  | Acute | Cell signalling |
| ***Mfap5*** | ↓ |  | Acute | ECM / fibrosis |
| ***Nppb*** |  | ↓ | Chronic | Cardiac stress marker |
| ***Nr1d1*** | ↓ |  | Acute | Nuclear receptor / metabolism |
| ***Nudt13*** | ↑ |  | Acute | Mitochondrial metabolism |
| ***Ogfr*** | ↑ |  | Acute | Cell growth regulation |
| ***Parp3*** | ↑ |  | Acute | DNA repair / modification |
| ***Pla2g16*** | ↑ |  | Acute | Lipid metabolism |
| ***Plgrkt*** | ↑ |  | Acute | Fibrinolysis / immune response |
| ***Pomp*** | ↑ |  | Acute | Proteasome maturation |
| ***Ppa1*** | ↑ |  | Acute | Energy metabolism |
| ***Psme1*** | ↑ |  | Acute | Proteasome activation |
| ***Rgcc*** | ↓ |  | Acute | Regulator of cell cycle |
| ***Rmdn3*** | ↑ |  | Acute | Regulator of microtubule dynamics |
| ***Rrad*** | ↑ |  | Acute | Cell signalling / GTPase |
| ***Saraf*** | ↑ |  | Acute | Calcium signalling |
| ***Sp100*** | ↑ |  | Acute | Immune response / nuclear antigen<br>Type I/II IFN-response |
| ***Stat1*** |  | ↑ | Chronic | Type I/II IFN signalling |
| ***Stat3*** | ↑ |  | Acute | Cell signalling / STAT pathway |
| ***Tapbp*** | ↑ |  | Acute | MHC I / antigen processing |
| ***Tcim*** | ↓ |  | Acute | Immune response |
| ***Tmem140*** | ↑ |  | Acute | Membrane protein / unknown function |
| ***Tnfrsf12a*** |  | ↓ | Chronic | TNF signalling / inflammation |
| ***Trafd1*** | ↑ |  | Acute | Immune signalling regulation<br>Type I/II IFN-response |
| ***Tyrobp*** | ↑ |  | Acute | Immune signalling adaptor |
| ***Uba7*** | ↑ |  | Acute | IFN-a/b induced ubiquitination |
| ***Vwa5a*** | ↑ |  | Acute | ECM / adhesion |
| **Unannotated transcript**<br>*(NewGene_537)* | ↑ |  | Acute | Unannotated transcript; function unknown |
| **Putative transcript**<br>*(NewGene_6563)* | ↑ |  | Acute | Putative MHC II / antigen presentation associated transcript |
| **Unannotated transcript**<br>*(NewGene_12)* |  | ↑ | Chronic | Putative MHC II / antigen presentation associated transcript |
| **Unannotated transcript**<br>*(NewGene_525)* |  | ↑ | Chronic | Putative IFN-induced secreted protein |

