## Supplementary figures and images for "Shared and divergent features of the immune response to acute and chronic *Trypanosoma cruzi* infection in mice revealed by cardiac transcriptomics"

### Supplementary Figure 1

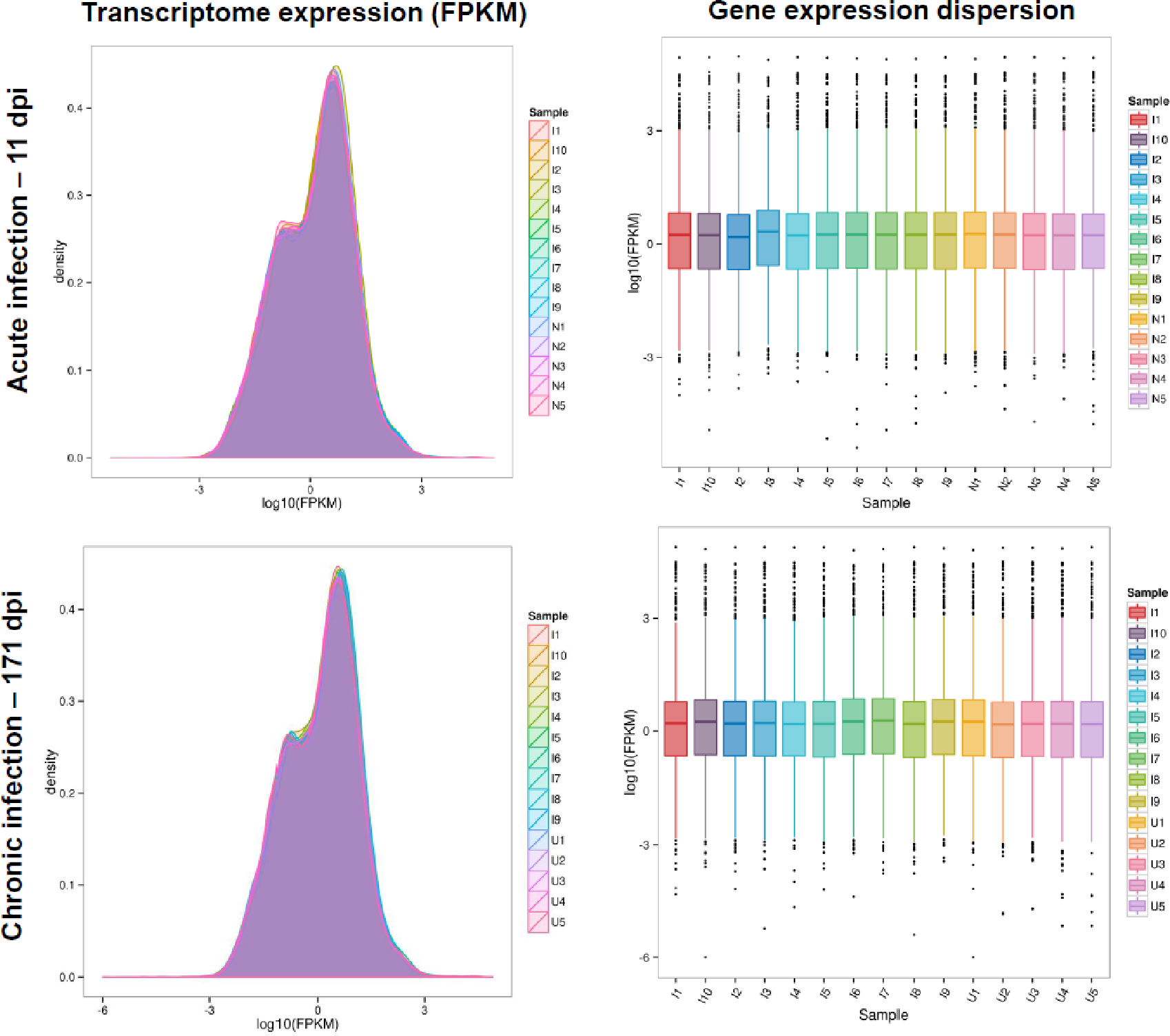

### Supplementary Figure 2

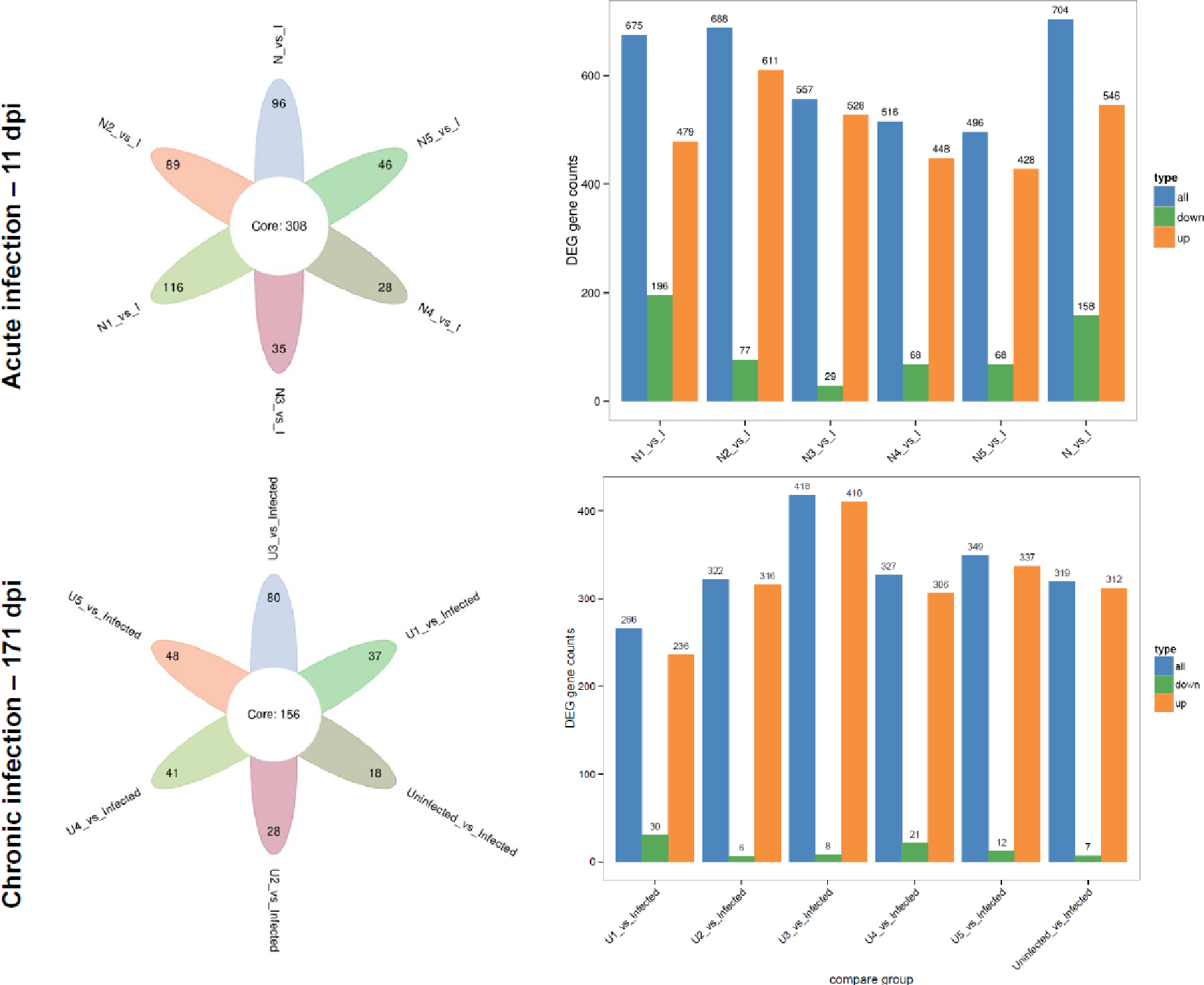

### Supplementary Figure 3

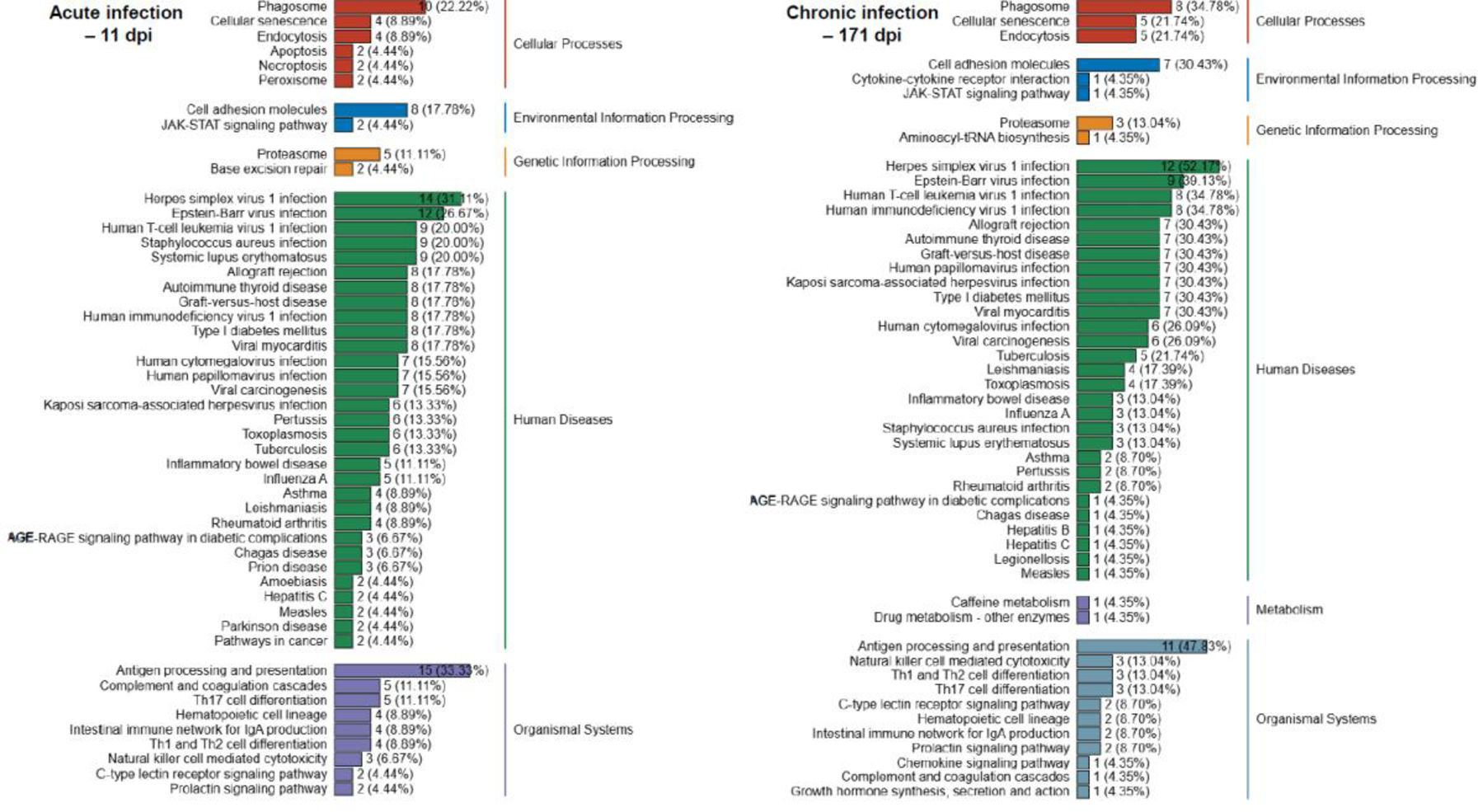

### Supplementary Figure 4

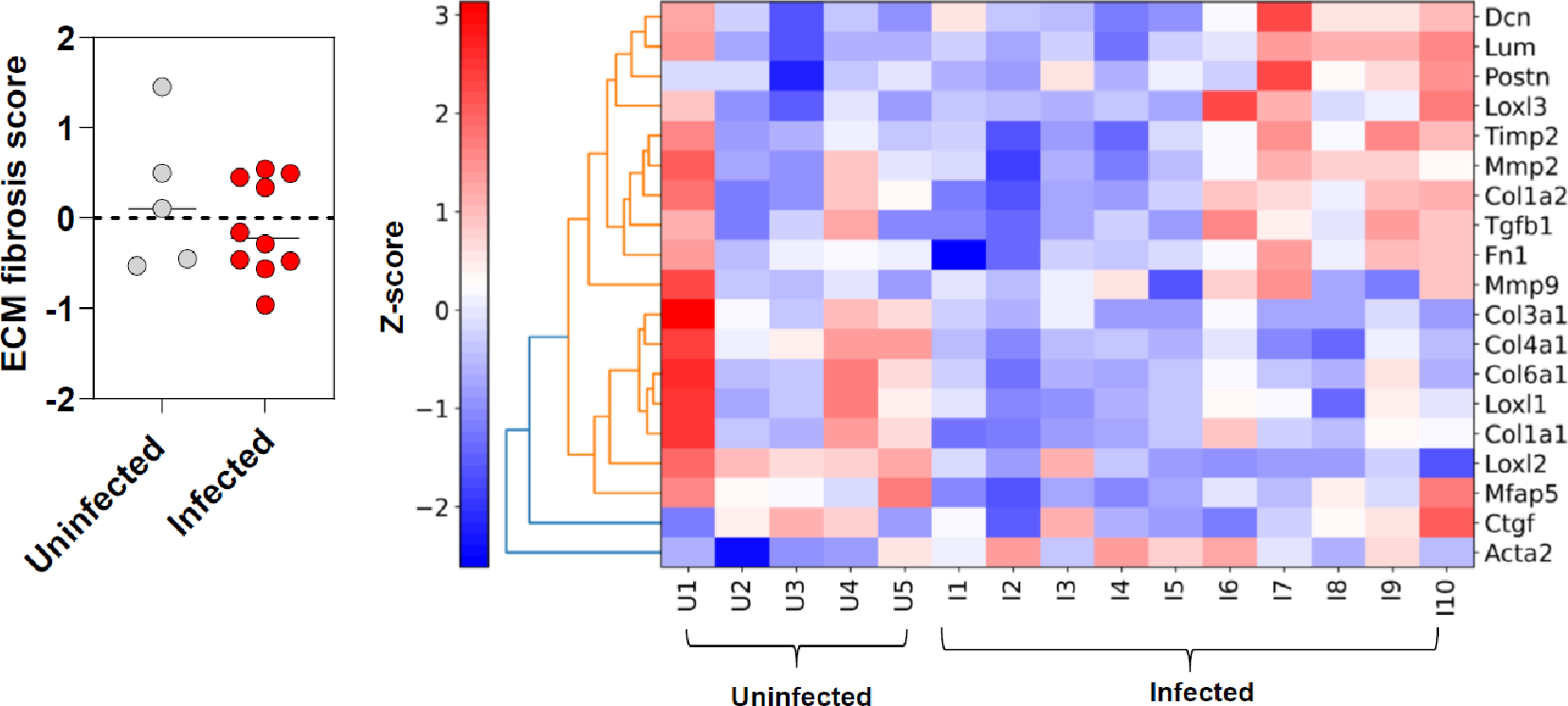
